# The geographic structure of chloroplast capture in a hybrid zone

**DOI:** 10.64898/2026.02.27.708378

**Authors:** N.J. Engle-Wrye, R.A. Folk

## Abstract

The frequency of hybridization differs strongly across different taxa and areas of Earth, yet the environmental drivers of this variation remain poorly understood. Areas strongly influenced by Pleistocene glaciation have long been hypothesized to be rich in hybrid taxa through forcing previously allopatric species into contact and interrupting ecological barriers. Here, we revisit naturally occurring *Heuchera americana* × *H. richardsonii* in North America using deep population level sampling of nuclear and plastid genomes to re-evaluate the long-standing hypothesis that Pleistocene climate dynamics promoted hybridization in this system. An extensive population-level sampling scheme involving nuclear and plastid genetic data (729 individuals across 455 populations) reveals a more complex hybridization history than implied by previous representative phylogenetic sampling. Our analyses identify multiple independent chloroplast capture events since diversification within this complex, rather than a previously hypothesized single ancestral event. Strong geographic structure at higher-level plastid clades but little structure within subclades, together with the rarity of ancestral plastid genomes, suggests a unidirectional inheritance of the introgressed plastid genome rather than a simple geographic distribution of plastid haplotypes. Ancestral niche projections indicate that Pleistocene glacial maxima created a shared narrow eastern refugium east of the Mississippi River, enabling localized hybridization, while western refugia were larger and taxa probably allopatric. This contrast led to an east-west gradient in cytonuclear discord. Our results corroborate and refine a 90-year-old hypothesis of climate-mediated hybridization by demonstrating that historical gene flow was spatially restricted but hybrid genotypes now command a larger geographic area. Our study highlights the importance of deep population-level sampling for accurately reconstructing hybridization histories and underscores the catalytic role of climatic instability in general as a driver of hybridization in *Heuchera*, and likely many other taxa across the tree of life.

## Introduction

Hybridization, the flow of genetic information between taxa, is one of the hallmarks of plant evolution, and in the form of cytonuclear discord it is both conspicuous and pervasive, with thousands of reports in the literature from the dawn of molecular systematics to the present (Rieseberg, 1991; Yang et al., 2023; Huang et al., 2025; Liu et al., 2025). Despite a deluge of research reporting cytonuclear discord in plants, its ecological drivers remain obscure, although geographic range disturbance is a leading mechanism (Anderson, 1949; Folk et al., 2018b, 2023). This knowledge gap is important because we know that the distribution of cytonuclear discord is highly heterogenous across the tree of life and geographic areas (Ellstrand et al., 1996), but we remain without generally agreed mechanisms for this disparity (Larson et al., 2026). Hence, while hybridization has been recognized as a major driver of novel variation and genomic datasets and coalescent methods have greatly increased the facility and pace of detecting it, studies of the principle drivers of cytonuclear discord and other forms of hybridization in plants lags behind other richly studied mechanisms such as polyploidy (Folk et al., 2018b).

The biological impact of Pleistocene glaciation, as a profound geological disruptor of geographic range, was profound and multidimensional, impacting species distributions (Birks, 2008), community assembly (Tóth et al., 2019), population genetics (Hofreiter and Stewart, 2009), and extinction dynamics (Faith and Surovell, 2009), ultimately resulting in a turnover of much of the world’s biota (Rabosky et al., 2018; Folk et al., 2019; Igea and Tanentzap, 2020), often in a life history-dependent manner (Lyons, 2003; Guralnick, 2007). The implication of this for hybridization was not lost on early investigators; a major outstanding hypothesis is that rapid geographic range shifts during the Pleistocene drove many of the hybridization patterns we see today (Anderson and Stebbins, 1954; Ikeda et al., 2012; Lopez-Alvarez et al., 2015; Marques et al., 2016; Folk et al., 2018b, c; Turchetto et al., 2022; Folk et al., 2023) via novel species contact patterns, the latter documented in numerous taxa by the fossil record and by paleoclimatic modeling/hindcasting (Roy et al., 1995; Graham et al., 1996; Lyons, 2003; Bush et al., 2004; deMenocal, 2004).

*Heuchera* (Saxifragaceae), a flowering plant clade characteristic of North American montane environments, is one of the most prolific examples in plants of interspecific hybridization, in terms of species number and evolutionary distance (see (Soltis et al., 1991; Soltis and Kuzoff, 1995; Folk et al., 2016). With a purported (Cronquist, 1947; Calder and Savile, 1959; Spongberg, 1972; Wilkins and Bohm, 1976) and demonstrated (Rosendahl et al., 1936; Wells, 1979) intrinsic propensity for hybridization, multiple evolutionarily replicated hybridization events in disparate clades (Folk et al., 2016), and occurrence across the wide environmental gradients of North America, *Heuchera* is an excellent evolutionary system for testing hypotheses about drivers of hybridization. Within this group, we chose the hybrid complex of *Heuchera americana*, *Heuchera richardsonii* and the named hybrid between them (*Heuchera americana* var. *hirsuticaulis*) as our specific study system to investigate Pleistocene glaciation as an abiotic driver of hybridization, expanding upon previous efforts that focus above the species level (Folk et al., 2018c, 2023), using a population sampling approach.

Rosendahl et al. (1933, 1936) hypothesized that before the southerly advance of the Pleistocene glaciers, *Heuchera richardsonii* and *H. americana* were allopatric species with strong geographic isolation but few intrinsic isolation mechanisms. They proposed that *H. americana* had a southeastern range, thought to center east of the Appalachians, while *H. richardsonii* had a northwestern range spanning the northern Great Plains to the western region of the American Midwest. In part because these plants are primarily endemic to rock outcrops, cliffs, and talus slopes (Folk et al., 2016), they are thought to have thrived on the expanding glacial margins to which they are well adapted, providing a potential mechanism promoting temporary sympatry (Folk et al., 2018b). As is common for closely related plant species (Anderson, 1949), disturbed sympatric distributions are thought to have spawned a hybrid zone, resulting in the hybrids currently taxonomically recognized as *Heuchera americana* var. *hirsuticaulis*. During a population rebound as the glaciers receded, Rosendahl et al. (1936) hypothesized that the resultant hybrid variety *Heuchera americana* var. *hirsuticaulis* completely replaced its parents over a large diagonal swathe across central North America from northern Louisiana to southern Michigan. Currently, *Heuchera richardsonii* is endemic primarily to the eastern Great Plains states with outliers in the western Great Plains north to the Canadian Interior Plains and Canadian Shield, while *H. americana* is primarily distributed in the eastern U.S. east of the Mississippi, with a disjunct population residing in the Ozarks and further outlier southerly populations to Texas. Intuitively, the distribution of the hybrid between these two species is restricted exclusively to a shared border region, in a broad band from southern Michigan to Louisiana, and completely replaces its parents, whose unhybridized forms never immediately co-occur either with each other or the hybrid (Wells, 1984).

While a compelling hypothesis, Rosendahl et al. (1936) relied on morphological data, afterwards corroborated by artificial crossing experiments demonstrating interfertility and morphologically similar plants to *H. americana* var. *hirsuticaulis*. The existence of ancient hybridization events in *Heuchera* was later corroborated by a phylogenomic study (Folk et al., 2016) that used genetic data from 42 of the 46 presently recognized *Heuchera* species; from which one *H. americana*, two *H. richardsonii*, and one hybrid were included. This and previous and subsequent work completing sampling of described taxa (Soltis et al., 1991; Soltis and Kuzoff, 1995; Okuyama et al., 2008; Folk et al., 2018c, 2023) suggests that the majority of *Heuchera* (25 of the 42 tested species) experienced cytonuclear discord, and (Folk et al., 2018c, 2023) pointed to Pleistocene-driven geographic range disturbance as the likely mechanism promoting overlap and hybridization. Pleistocene disturbance as a driver is attractive as it provides a ready explanation for otherwise illogical allopatry, polyphyly, or other disparities across geographic areas and taxa.

The hypothesis of Folk et al. (2016) for the *H. americana* clade was that “plastid DNA introgressed into a single ancestral eastern species of section *Heuchera* from one of several sympatric members of section *Holochloa*” on the basis of a monophyletic clade exclusive of *H. richardsonii* being monomorphic for a captured chloroplast (“clade A” in that study, a clade that is not directly related to other *Heuchera* plastid clades). Preliminary results prepared for this study from further western population samples violated this monomorphic plastid state, suggesting a more complex chloroplast capture scenario in need of further investigation as it would likely revise the origin of the *H. americana × H. richardsonii* hybrid zone. Here, we aim to expand phylogenetic sampling to further test this hypothesis, pairing deep phylogenomic sampling with niche modeling to investigate in detail the potential role of historical climate change in these chloroplast capture events at a lower phylogenetic level and assess the overlap between refugia and the current distribution of plastid genomes. We hypothesize that (1) Pleistocene glacial maxima forced the suitable habitat of previously allopatric but closely related species *Heuchera americana* and *Heuchera richardsonii* into at least one shared refugium where contact and hybridization occurred and that 2) increased population-level sampling would reveal spatial patterns of hybridization and allow for the biogeographic inference of where hybridization occurred.

## Methods

### Sampling

We structured our population level sampling to cover species ranges with representative sampling of morphotypes corresponding to varieties delimited in Rosendahl et al. (1936). While subspecific taxa are not currently accepted in the hybrid or non-hybrid populations of either parental species, we aimed to investigate whether there was a correlation between these morphotypes and hybrid ancestry. Counted among *H. americana* var. *americana, H. richardsonii,* and *H. americana* var. *hirsuticaulis* as currently recognized are the following unrecognized varieties: *H. americana* var. *calycosa*, *H. americana* var. *brevipetala*, *H. americana* var. *americana* (syn. *H. americana* var. *typica*), *H. americana* var. *interior*, *H. richardsonii* var. *affinis*, *H. richarsonii* var. *grayana,* and *H. richardsonii* var. *typica*. All previous sampling efforts focused on *H. americana* (Folk et al. 2016; Folk et al. 2023) included only samples to the east of the hybrid’s range east of the Mississippi River, where the vast majority of the range lay. Importantly, we were aware of a few previously unsampled *H. americana* populations residing to the west of the hybrid zone, of rare occurrence across the Ozark periphery of Arkansas, north Louisiana, and east Texas, and included them to assess the status of these outlier populations, some of which have only recently been clarified (Holmes et al., 2011; Kelley, 2021). In total, we sampled 729 in- and outgroup individuals from 350 ingroup and 105 outgroup populations; averaging ∼1.6 individuals collected from each population. These collections were obtained through both field collections (up to 10 individuals per population) and through destructive sampling of herbarium specimens (1 individual per population due to the limitations of identifying unique genets from specimens). Unique population totals were 235 *H. americana* var. *americana*, 69 *H. americana* var. *hirsuticaulis*, 46 *H. richardsonii* populations, and 105 outgroup *Heuchera*; while unique individuals totaled 354 *H. americana* var. *americana*, 147 *H. americana* var. *hirsuticaulis*, and 102 *H. richardsonii* populations, and 126 outgroup *Heuchera*; (https://github.com/njenglewrye/The-geographic-structure-of-chloroplast-capture-in-a-hybrid-zone-manuscript/tree/main). Extensive outgroup sampling was warranted because (1) previous investigation suggests that *H. americana* is paraphyletic (Folk et al., 2018a); (2) *H. americana* and *H. richardsonii* are maximally distant in the clade *Heuchera* subsect. *Heuchera,* thus representatives of all members of *Heuchera* section *Heuchera* should be sampled; and (3) analysis of *Heuchera* cytoplasmic DNA recovers three distant clades separated by numerous outgroups (Folk et al. 2016, 2018c, 2023).

### Phylogenetic construction

DNA extraction used a modified version of CTAB DNA extraction approach (Doyle and Doyle, 1987) with modifications described in (Folk and Freudenstein, 2014). Illumina libraries were built using NEBNext Ultra II prep kits. We performed sequence capture with biotinylated RNA baits that target a customized panel of 277 low-copy nuclear loci specific to *Heuchera* (details following Folk et al. 2015), but also yields high amounts of off target cpDNA and mtDNA (Folk et al., 2025). Finally, we sequenced the captured libraries on an Illumina Hiseq 3000 instrument (Novogene, San Diego, California).

Plastid alignments followed methods established in Folk et al. (2016). Read pileups were made using BWA-MEM (Li, 2013) against the *H. parviflora* var. *saurensis* plastid genome (GenBank ID: KR478645) followed by variant calling and consensus sequence generation in SAMtools, bcftools and vcfutils.pl (Danecek et al., 2021); data were treated as haploid. The plastid assembly was trimmed for missing data by removing nucleotide sites with greater than 50% missing data and taxa with greater than 20% missing data. A phylogeny was then inferred using a concatenated analysis in RAxML-NG (Kozlov et al., 2019) with a GTR+Γ model with genes treated as distinct partitions and the tree was rooted on *Heuchera elegans* (“clade C” following (Okuyama et al., 2005; Folk et al., 2016), likely ultimately from *Mitella* and absent in the study region). Two further clades were identified in the phylogenetic inference: “clade B,” a clade with no immediate relatives considered by Folk et al. (2016) and Soltis et al. (1991) to be the ancestral plastid type, and clade A, considered by Folk et al. and Soltis et al. to represent introgression, likely ultimately from an ancestor of *Elmera* (Folk et al. 2023). To visualize whether patterns of chloroplast capture were regional, we geographically mapped the *Heuchera* populations and color coded them to phylogenetic placement, both highlighting the distribution of clades A and B and highlighting major subclades of CP capture clade A. Finally, a tanglegram was constructed using the nuclear ASTRAL tree from BIORXIV/2026/708067 versus our RAxML plastid phylogeny, using phytools in R Studio Version 2023.03.1+446 (Revell, 2024).

### Dating

To infer a timeline for ancestral niche reconstruction, we repeated the time calibration strategy of Folk et al. (2023), a dating analysis across the broader tribe Heuchereae. This analysis was modified by adding an accession of *H. americana* var. *hirsuticaulis* to calibrate this additional internal node and completely represent the three ingroup taxa. Other details follow Folk et. al (2023) and briefly comprised a concatenated analysis of the nuclear gene set implemented in MCMCtree (Yang and Rannala, 2006; Yang, 2007) with three node calibrations on the stem of *Heuchera* and two other locations in tribe Heuchereae based on (Deng et al., 2015; Folk et al., 2023).

### Occurrence records and distribution modeling

Occurrence records were drawn from verified records in Folk et al. (2023) with the exception that records of *H. americana* var. *americana* and *H. americana* var. *hirsuticaulis* were split on a geographic basis (following Wells, 1984) for the purpose of this analysis. Models were reinferred according to the parameters of the earlier analysis in Maxent with identical training set and bootstrap parameters, which are outlined in Folk et al. (2023).

### Ancestral niche reconstruction

We first determined environmental tolerances of extant species by making species distribution models using MaxEnt (Phillips et al., 2017), following the same parameters as (Folk et al., 2018c, 2023). This analysis occurs in two stages, namely (1) ancestral reconstruction of niche parameters and (2) projection of these parameters onto climate scenarios. Step (1) was performed across tribe Heuchereae so as to provide full outgroup context for ancestral reconstruction, while step (2) was narrowed down to the nodes representing the focal three taxa as outgroups were not relevant for this purpose. The twelve environmental predictors used followed (Folk et al., 2018c, 2019, 2023) and included: four Bioclim variables (Hijmans et al., 2005): of mean annual temperature (Bio1), temperature annual range (Bio7), annual precipitation (Bio12), and precipitation of the driest quarter (Bio17); two topographical variables of elevation and slope (https://doi.org/10.5066/F7DF6PQS); four SoilGrids250m variables (Hengl et al., 2017) of mean coarse fragment percentage, mean pH, mean sand percentage, and mean organic carbon content; finally, our two land cover variables were needle-leaf and herbaceous land cover (Tuanmu and Jetz, 2014). From the averaged models, a custom Predicted Niche Occupancy (PNO) calculator (https://github.com/ryanafolk/pno_calc (Folk et al., 2023)) was used to extract climatic data for ancestral niche reconstruction for all taxa but *H*. *americana* var. *hispida*, for which we directly extracted environmental data from the occurrence points due to this taxon having fewer verified records. Then we procured historical layers, for mean annual temperature (Bio1) and precipitation of the driest quarter (Bio17), using PaleoClim (Brown et al., 2018) and used Utremi to model probabilities of suitable habitat through the 51 time points of (Folk et al., 2023) from 0 to 3.3 MYA. Previous work (Folk et al., 2023; Dahal et al., 2025) and preliminary analyses (see supplementary figure B.3) with Utremi demonstrated that Bio1 primarily provides habitat suitability information on a north to south axis (i.e., primarily temperature clines), but we wished to also constrain the ancestral projection in the east to west axis as it is unlikely western suitable habitat was dispersible given the extant range of these taxa. For this reason, we included Bio17 (precipitation of driest quarter, a measure of seasonality as well as absolute precipitation). Utremi was originally designed for univariate analyses but it was straightforward to implement a multivariate analysis through multiplying site probabilities (i.e., joint probability of suitable habitat) between Bio1 and Bio17 after the final projection step.

## Results

### Phylogenetics and chloroplast capture

As repeatedly recovered previously (Soltis et al. 1991; Okuyama et al. 2005; Folk et al. 2016; Folk et al. 2023), *Heuchera* was recovered as polyphyletic with species distributed in either ancestral plastid clade B (i.e., the clade considered to represent those taxa lacking chloroplast capture; Folk et al. 2016) or plastid clade A; see Fig. 4.1 inset (detailed view with individual tip labels in Supplementary Fig. B.1). Ancestral plastid clade B contained a minority of taxa in the study group: 37 tips with six *H. americana* var. *americana*, two *H. americana* var. *hirsuticaulis*, 18 *H. richardsonii*, and 11 outgroup *Heuchera* previously recovered in this clade. Chloroplast capture clade A had 691 tips (i.e., by far the majority of accessions in the study group), with 348 *H. americana* var. *americana*, 145 *H. americana* var. *hirsuticaulis*, 84 *H. richardsonii*, and 114 outgroup *Heuchera* (notably, all 45 accessions of *H. americana* var. *hispida*, a more eastern hybrid between *H. americana* and *H. pubescens*, were placed here but are not considered further below). The tanglegram between the plastid and nuclear phylogenies (Fig. B.2) showed high levels of discordance among the evolutionary histories of nuclear and plastid genomes in almost all of the *Heuchera* included in this study, which was similar to Folk et al (2016). Essentially, the two genomes completely lack a shared phylogenetic backbone in any respect. We prepared color-coded maps denoting plastid clade memberships (Fig. 4.1), which showed highly distinct geographic regions for populations belonging to either capture clade A or ancestral clade B (Fig. 4.1A), but little geographic pattern among the main highly supported subclades of CP capture clade A (Fig. 4.1B). Thus higher-level plastid clades possess strong geographic signal whereas lower-level clades do not. Most notably, Fig. 4.1A contains an unexpected result that contradicts Folk et al. (2016): both parental taxa contain both plastid clades A and B. B-containing (likely ancestral) haplotypes are restricted to west of the Mississippi River without exception and occupy this region to the exclusion of the neighborhood of *H. americana* var. *hirsuticaulis*.

### Ancestral niche reconstruction

Ancestral projection in Utremi inferred niche probability maps that illustrate migration of refugial habitats. From the 35 maps, we selected six characteristic maps representing glacial minima and maxima and their corresponding typical habitat suitabilities, shown Fig. 4.2 (a complete animated GIF of the geographic projection is available at https://github.com/njenglewrye/The-geographic-structure-of-chloroplast-capture-in-a-hybrid-zone-manuscript/blob/f938a2adc344218af3a85429dbd3fc9be6990499/Niche%20reconstructions/Bio1XBio17/combinedBio1Bio17_normal.gif). Geographic projection identified three discrete regions of habitat suitability, from west to east referred to as the western, central, and eastern refugia. These refugia (which represent refugial habitat suitability rather than dispersal processes given the limitations of the model) were reconstructed at their widest distributions during glacial minima and as such we describe them in the context of refugia modeled at 2.51 mya (approximately the Plio-Pleistocene boundary), Fig. 4.2. The western refugium ranges from California to west Texas and from Canada down into Mexico; this climate refugium is very distant from other refugia and the study group was considered unlikely to have dispersed there given the solely eastern-central distribution of *Heuchera* subsect. *Heuchera* (its sister clade *H.* subsect. *Parvifoliae* instead occupies this region); thus this climate refugium is not considered further. The central refugium ranges from western Oklahoma to Minnesota and Wisconsin, corresponding closely to the modern range of *Heuchera richardsonii*. The eastern refugium ranges diagonally from Arkansas to Maine and down to Georgia, closely resembling the modern range of *Heuchera americana* var. *americana*. At 2.51 mya, there was no appreciable habitat suitability in the modern range of *H. americana* var. *hirsuticaulis*, suggesting allopatry. During later glacial maxima (e.g., 1.58 mya; Fig. 4.2), the refugial reconstructions are less discernable with smaller southerly disjunctions (particularly gaps in the floodplain of the Mississippi and Apalachicola rivers). The suitability in this timeframe ranges from Texas down into Mexico, and the prediction is generally more westerly than the modern distribution of the study group, but sometimes as far east as Georgia. Floodplain gaps were also observed using only Bio1 (Supplemental Fig. 4.3), similar to the modern distribution. The eastern climate refugium drops out entirely during certain glacial minima, e.g., 1.12 mya, or overlaps with other refugia in southern Georgia and Alabama, as seen 1.58 mya (Fig. 4.2), thus indicating a range constriction or microrefugia.

## Discussion

### Ancestral niche reconstructions

Ancestral niche reconstructions indicate that warm late-Pleistocene periods were associated with two broad ancestral regions of suitable habitat similar to contemporary species distributions, which were interrupted by glacial maxima. Suitable habitat at glacial maxima predicts a pair of greatly reduced southerly refugia that would have resulted in range overlap among parental species. Among these, the western refugium was consistently more widespread, while the eastern refugium was often so narrow our model showed the near-disappearance of suitable habitat, pointing to a role for refugial microhabitats. This refugial reconstruction of a west-to-east increase in range disturbance leads to the prediction that easterly populations experience greater cytonuclear discord. Accordingly, we recovered a strong west-to-east pattern of plastid clade membership. All populations of *Heuchera americana* var. *americana* and *H. richardsonii* bearing the ancestral B plastid clade are west of the Mississippi river. Chloroplast capture clade A occurs without exception east of the Mississippi River. In a corridor from the Ozarks to the Great Lakes region, chloroplast capture clade A occupies the range of the hybrid *H. americana* var. *hirsuticaulis* (which is monomorphic for this plastid) and neighboring populations of *H. richardsonii*. These geographic areas correspond remarkably well to the original morphological hypothesis (Rosendahl et al., 1936).

### Revision of chloroplast capture

The most important overall finding was evidence for a more complex hybridization scenario. Folk et al. (2016), which sampled one accession per taxon, suggested that a single chloroplast capture event occurred in the ancestor of the clade comprising all *Heuchera* subsect. *Heuchera* except *H. richardsonii* on the basis of plastid clade monomorphy evident at that time. Increasing sampling depth to the population level, has falsified this previous hypothesis of a single chloroplast capture event because *H. americana* var. *americana* and *H. richardsonii* contain both plastid genome types. This revised clade distribution requires at least three chloroplast capture events in the history of the hybrid complex, at least two involving eastern members of section *Heuchera* exclusive of *H. richardsonii* (so as to account for the presence of plastid clade B in extreme westerly populations of *H. americana* var. *americana*, but its lack in every remaining taxon in *H.* subsect. *Heuchera*), and a third event involving easterly populations of *H. richardsonii*.

Population mapping of plastid clade memberships (Fig. 4.1) appears to imply that plastid clade A inheritance was unidirectional. This is suggested because without exception, all included populations east of the Mississippi River are claded in clade A, leaving a disjunct plastid clade B distribution. Membership in plastid clade B is found in two disjunct areas, comprising most populations of *H. americana* var. *americana* in and below the southern Ozarks, and most populations of *H. richardsonii* west of the Great Lakes region. Aside from geographic distribution, it is also remarkable that only 4.3% (26/603) of the focal taxa placed in plastid clade B. The imbalanced geographic and phylogenetic distribution indicates the asymmetric nature of hybridization. This geographic pattern could be consistent with selection, namely cyto-nuclear incompatibilities induced in hybrids with the ancestral clade B plastid, or selective advantages of plastid clade A in a hybrid background (see review in Sloan et al. 2017). While determination of the mechanism likely requires experimental evidence, these results indicate chloroplast capture clade A may be spreading geographically, concordant with the complete lack of spatial structure in its subclades (Fig. 4.1B). Membership in clade B in two disjunct areas is best explained as a remnant of an earlier geographic structuring extant due to differing and separate refugia in westerly areas. Overall, the data suggest a refugium east of the Mississippi River where hybridization occurred, and refugia west of the Mississippi River in which hybridization did not occur or was infrequent. The distribution of plastid clade membership does not correspond to any previously recognized subspecific taxon. Likewise, color coding subclades of CP capture clade A (Fig. 4.1B) revealed that there was little to no geographic structure within the chloroplast capture clade, which would correspond to these taxa, most of which occur east of the Mississippi River.

### Pleistocene-driven hybridization

We find strong evidence for and expand upon Rosendahl et al. (1936) hypothesis of climatically mediated hybridization in *Heuchera*. Phylogenetic data, ancestral niche reconstructions, and contemporary species distributions all align well with the original Rosendahl et al. (1936) hypothesis that Pleistocene glacial mediated distributional change was the primary driver of hybridization between *H. richardsonii* and *H. americana*. Rosendahl et al. based their inference solely on morphological and biogeographic data, particularly the contrast between *H. richardsonii* solely occupying formerly glaciated areas and *H. americana* var. *americana* occupying areas south of maximal glaciation, which they therefore believed to be ancestrally allopatric, while *H. americana* var. *hirsuticaulis* occupies the contact zone between these two taxa. Our study as well as Folk et al. (2023) are consistent with this verbal model in that our Utremi analysis reconstructs maximal range overlap of hybrid parents during glacial maxima of the Pleistocene period, whereas earlier Pliocene reconstructions indicate a disjunct occurrence.

Our Utremi results add information on timing the hybridization event. We inferred disjunct suitable habitats around 3.32 mya and 2.51 mya (Fig.4.3) and less afterward, pointing to a Pleistocene date for hybridization. Ranges overlapped at glacial maxima, around 1.58 mya and 1.12 mya (Fig. 4.2); thus range overlap appears to localize near the mid-Pleistocene boundary when glaciation became more extreme. However, our study significantly extends previous inferences by demonstrating that contact and hybridization was not widespread across all contact zones but appears to have been confined to an eastern refugial zone. Though there were likely populations of both parental taxa in the glacial refugium spanning Texas and Louisiana, they were probably more dispersed and hybridized less frequently among larger populations, as they are modeled to have had broader suitable habitats 1.58 mya and 1.12 mya. By contrast, the refugium to the east of the Mississippi River had less reconstructed suitable habitat, pointing to reduced population sizes for the taxa in the eastern refugium. The latter scenario would be consistent with a population bottleneck and limited mate availability, both conditions consistent with the rapid expansion of chloroplast capture clade A among eastern members of the *H. americana* × *H. richardsonii* complex.

Utremi enabled us to model and visualize probabilities of historically suitable habitat, but it lacks a dispersal component to directly model the movement of the taxa, and thus should be seen as indicating climatic refugia rather than directly reconstructing distributional refugia. An additional limitation, shared with other ancestral reconstruction methods, is that reconstruction is performed at the level of species and does not explicitly consider population structure.

## Conclusions

This work emphasizes broad population-level sampling, rather than conventional macroevolutionary sampling practice, for accurately inferring historical patterns of gene flow between taxa. It will be recalled that *Heuchera americana* var. *americana* × *H. richardsonii* is in part a phylogenetic problem as the parents are non-sister. It has been asserted previously that population sampling may be ignored in practical situations for hybridization inference (Hibbins and Hahn, 2022). The necessity of taxon sampling has been recognized for decades for inferring organismal history generally (e.g., for correct inference of the flowering plant backbone; Soltis and Soltis, 2004), yet ancient hybridization tends to be perceived as a phylogenetic problem rather than a population genetics problem as in Hibbins and Hahn (2021); our work appears to be the first empirical interrogation of prevalent sampling practice for this application. Regardless of timeline, hybridization remains a population-level process and prevalent single-accession taxon sampling thus should systematically underestimate the complexity of hybridization. Thoughtfully constructed sampling across the geographic ranges of focal taxa can capture localized histories that might otherwise go undetected.

Expanding our population sampling has falsified the previous simplistic scenario of a single chloroplast capture event in the *H. americana* clade, revealing that rare *H. americana* populations in the western refugial area (west of the Mississippi river) retain the ancestral plastid but easterly *H. richardsonii* possess the derived plastid type, and therefore multiple interspecific chloroplast capture events are needed. Incomplete sampling in the prior therefore remarkably underestimated divergence among plastid haplotypes within both species by several million years. Identifying the geographic range of genetically pure cytoplasms of both parental species independently corroborates the finding from ancestral niche modeling that hybridization and chloroplast capture likely happened in the refugium east of the Mississippi river but not in the larger western refugium. Deeper sampling reveals that hybridization events are more numerous and localized than previously recognized. Our results found support for a classic hypothesis of Pleistocene-mediated gene flow among these two parental species, and more generally point to Pleistocene-influenced landscapes as cytonuclear discord hotspots.

## 1.1. Figures

**Figure 1.1.**
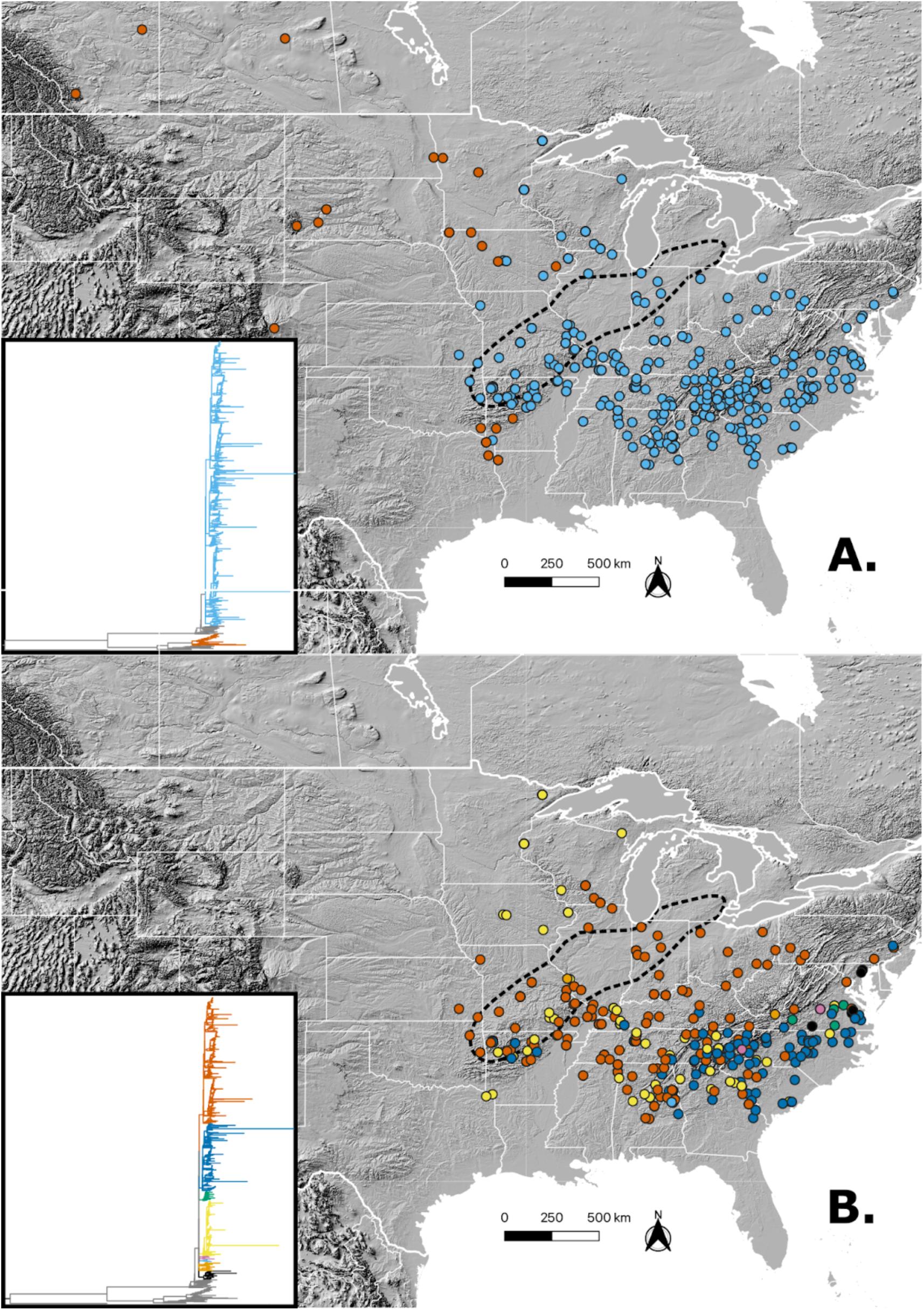
Color coded plastid clade phylogenies and maps (A) Plastid clade phylogeny and map with chloroplast capture clade A in blue and ancestral plastid clade B in orange, with the overall phylogenetic topology indicated in the left inset. (B) Map showing the distribution of subclades only of chloroplast capture clade A, with colors of populations on the map matching the colors of subclades in the phylogeny legend in left inset. In both (A) and (B), the dotted oval indicates the distribution of *H. americana* var. *hirsuticaulis* following Rosendahl et al (1936). All individuals southeast of this range are taxonomically treated as *H. americana* var. *americana*, and all northeast as *H. richardsonii*.

**Figure 1.2.**
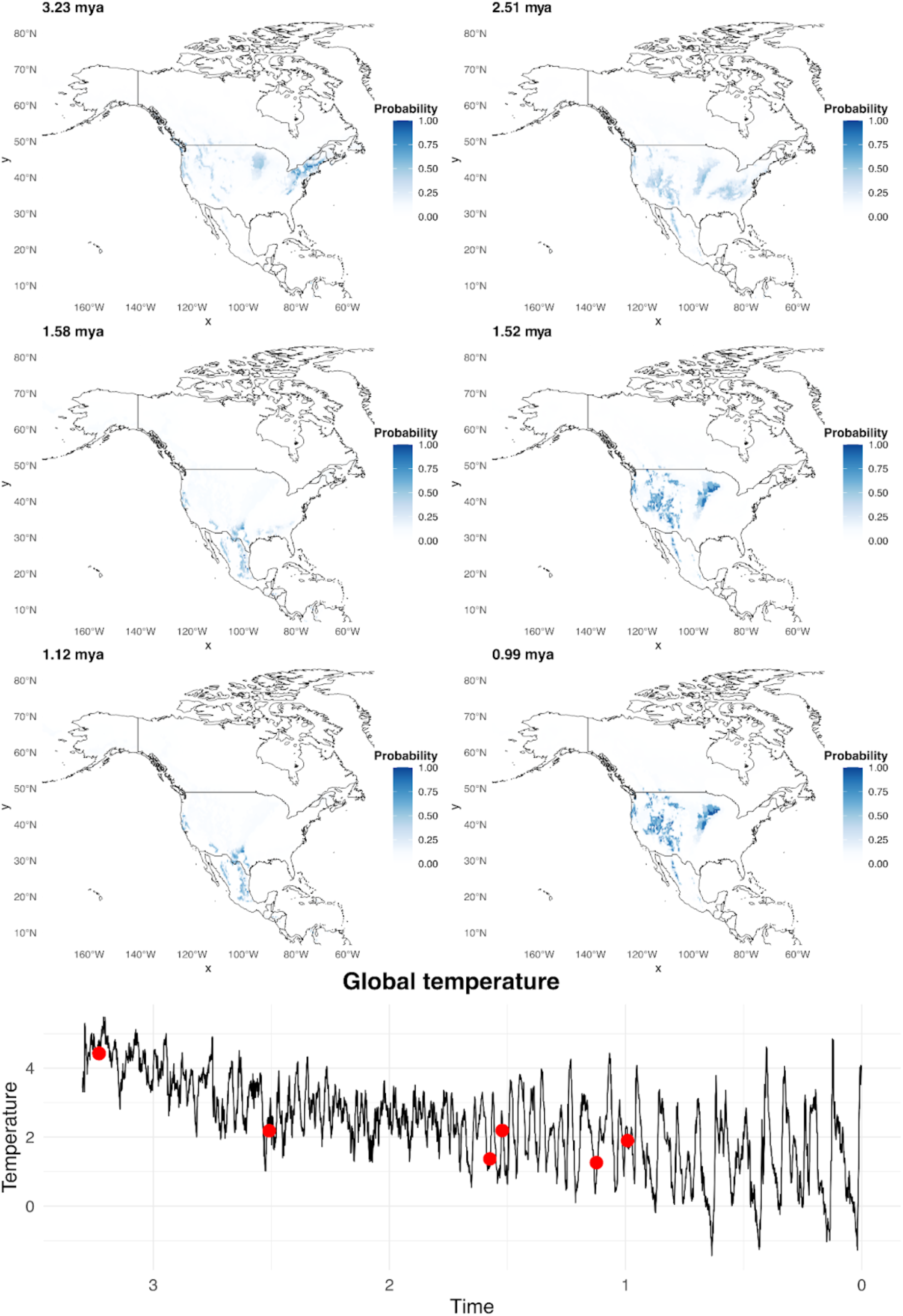
Utremi inferred habitat suitabilities. Maps show probabilities of suitable habitat matching the legends on their right, at various time points from 3.23 mya to 0.99 mya, inferred via Utremi using mean annual temperature (BIO1) and precipitation of the driest quarter (BIO17). Each map is represented by a red dot on the bottom line graph that shows time on the x-axis and temperature on the y-axis. Note that during early warmer periods there was a widespread refugium, likely only containing *Heuchera americana*, over the eastern states; and there was a second refugium over the midwestern states, likely only containing *Heuchera richardsonii*. Although there appears to be a third scattered range of suitable habitat amongst the far western states, we find it unlikely that these taxa were in those regions at these timepoints; the sister of *H. subsect. Heuchera* currently occupies this range. At 1.58 mya we see potential for sympatric species distributions and in the following timepoints we see the disappearance or extreme limitation of suitable habitat in the east. We think the latter is the case and represents a bottleneck that enabled the chloroplast capture found in the eastern *Heuchera* of this study. See Supplemental Figure 3 which represents Utremi results using only BIO1 and shows two clear refugia separated by the Mississippi river basin

## Acknowledgements

We would like to thank the curators at MEM, TEX, UNA, and NCU for allowing us to visit and collect specimens. Additionally, we would like to thank the NSF CAREER program (NSF DEB-2337784) and the Mississippi State University Biology Faculty Fund Research Award (Spring 2020) for funding.

## Notes

https://github.com/njenglewrye/The-geographic-structure-of-chloroplast-capture-in-a-hybrid-zone-manuscript/blob/f938a2adc344218af3a85429dbd3fc9be6990499/Niche%20reconstructions/Bio1XBio17/combinedBio1Bio17_normal.gif

https://github.com/ryanafolk/pno_calc

https://github.com/njenglewrye/The-geographic-structure-of-chloroplast-capture-in-a-hybrid-zone-manuscript/tree/main

### Competing Interest Statement

The authors have declared no competing interest.

