## Supplemental figures for "The geographic structure of chloroplast capture in a hybrid zone"

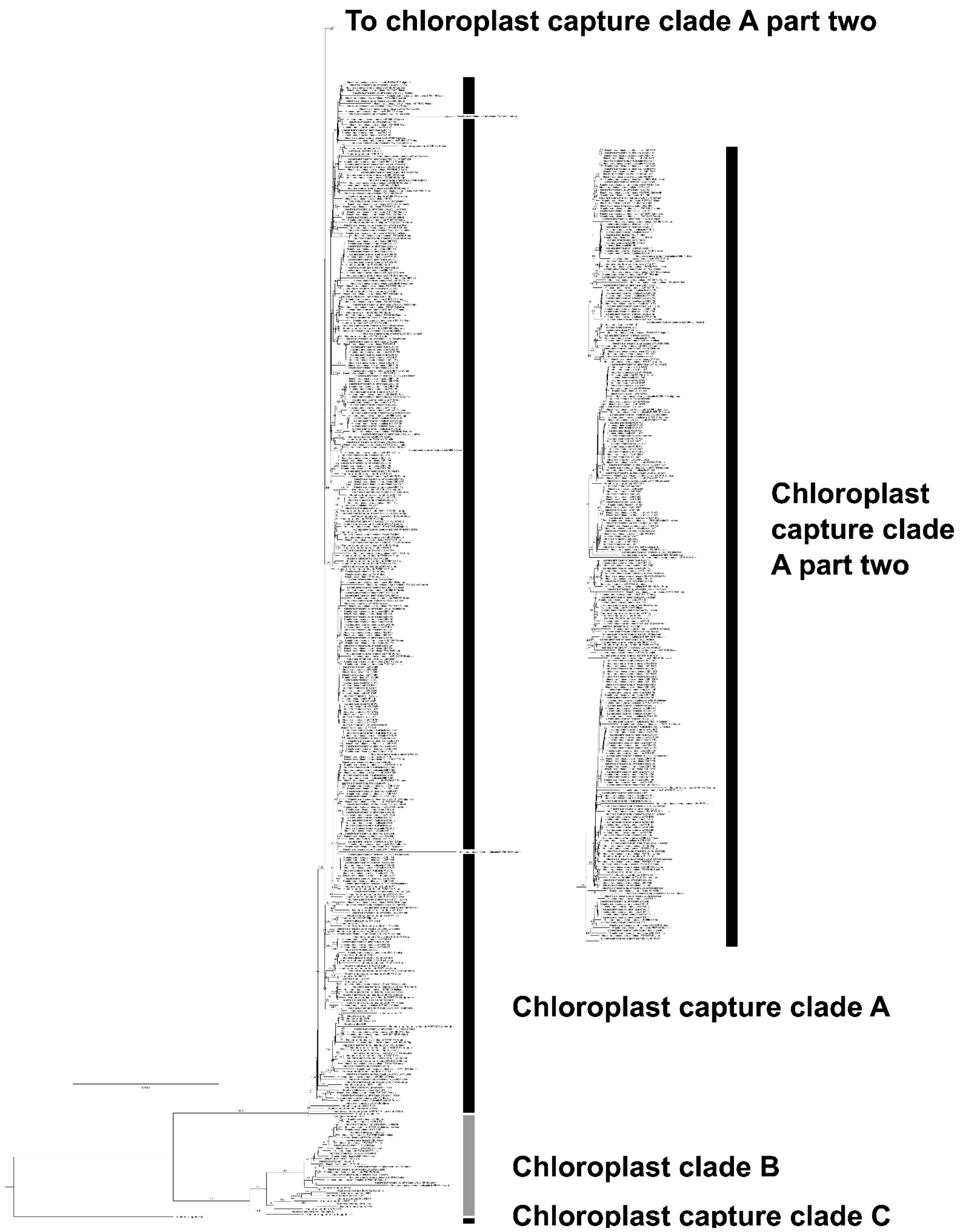


**Fig. S1 |** Chloroplast phylogeny with chloroplast clades labeled with a bar corresponding to the three clades: chloroplast capture clade C at the root, ancestral chloroplast clade B as the next diverging clade, and the rest of the phylogeny belonging to chloroplast capture clade A.

Chloroplast clade A has been clipped and continued to the right to save space.

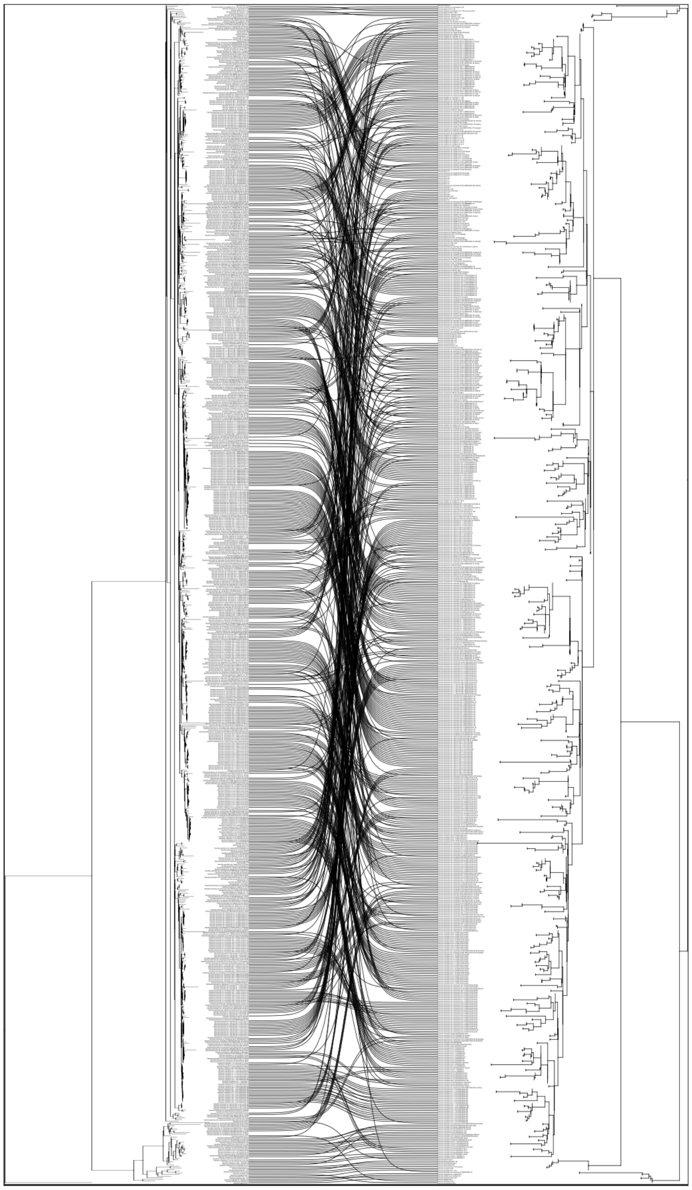

**Fig. S2 |** Chloroplast (left) versus nuclear (right) tanglegram. Lines connect shared accessions.

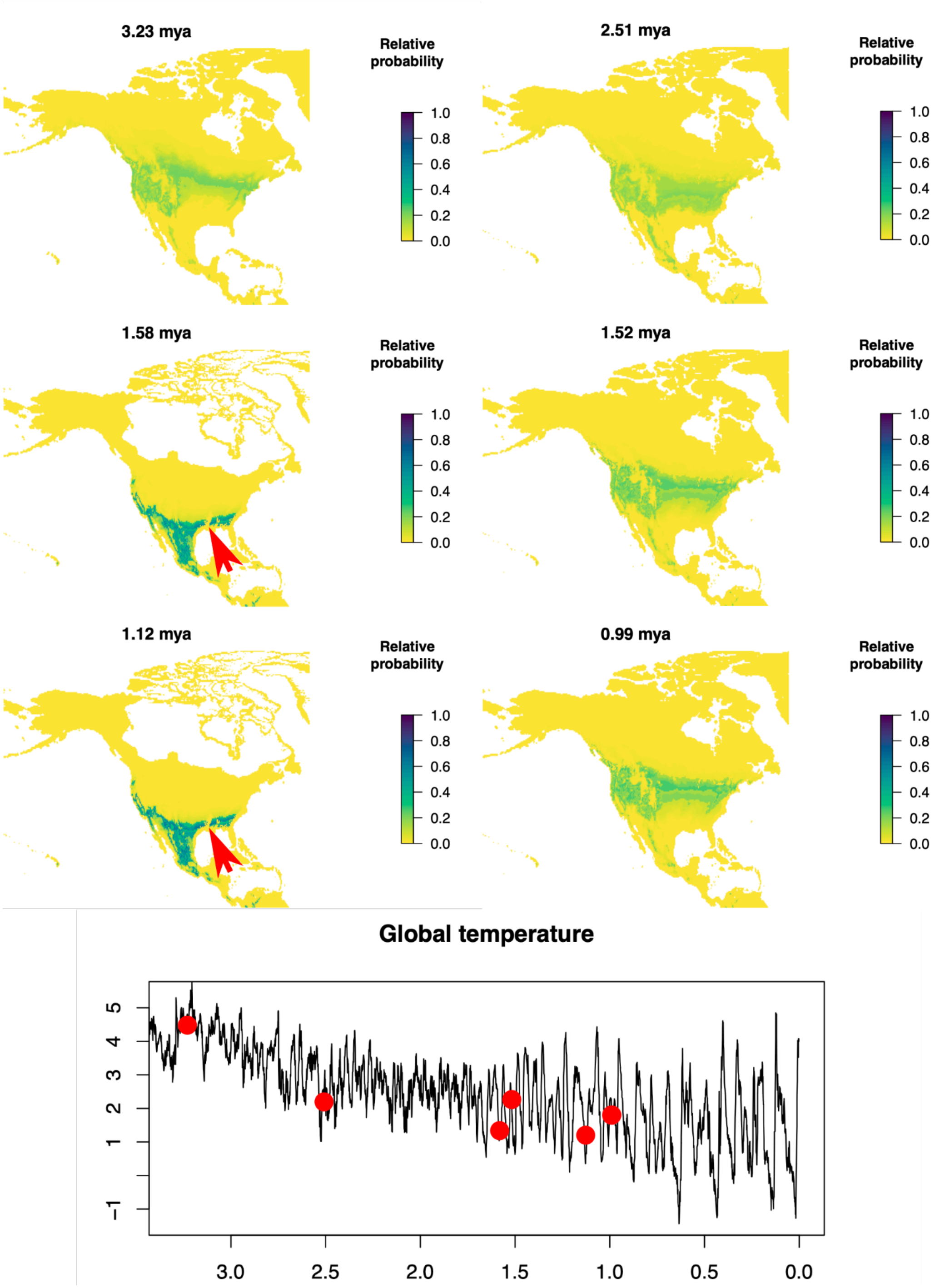


**Fig. S3 |** Utremi results using only mean annual temperature (BIO1) show a broad high probability band of habitat suitability from the Pacific to Atlantic coast with a gap between refugium at the Mississippi river basin, indicated by a red arrow.
